# *In Vivo* Biocompatibility of ZIF-8 for Slow Release via Intranasal Administration

**DOI:** 10.1101/2023.01.07.523104

**Authors:** Sneha Kumari, Thomas S. Howlett, Ryanne N. Ehrman, Shailendra Koirala, Orikeda Trashi, Ikeda Trashi, Yalini H. Wijesundara, Jeremiah J. Gassensmith

**Affiliations:** Department of Chemistry and Biochemistry, The University of Texas at Dallas, Richardson, TX, 75080; Department of Biomedical Engineering, The University of Texas at Dallas, Richardson, TX, 75080

**Keywords:** slow release, zeolitic imidazolate frameworks, depot effect, intranasal delivery

## Abstract

Zeolitic Imidazolate Framework-8 (ZIF-8) is becoming popular in research for its potential in antigen protection and for providing a thermally stable, slow-release platform. While papers applying these materials for immunological applications are aplenty in literature, studies that explore the biosafety of ZIF-8 in mammals—especially when administered intranasally—are not well represented. We checked the body clearance of uncoated and ZIF-coated liposomes and observed that the release slowed as ZIF-8 is easily degraded by mucosal fluid in the nasal cavity. We delivered varying doses of ZIF-8, checked their short- and long-term effects on diagnostic proteins found in blood serum, and found no noticeable differences from the saline control group. We also studied their lung diffusing capacity and tissue morphology; neither showed significant changes in morphology or function.

Graphical Abstract:
General overview of the investigation

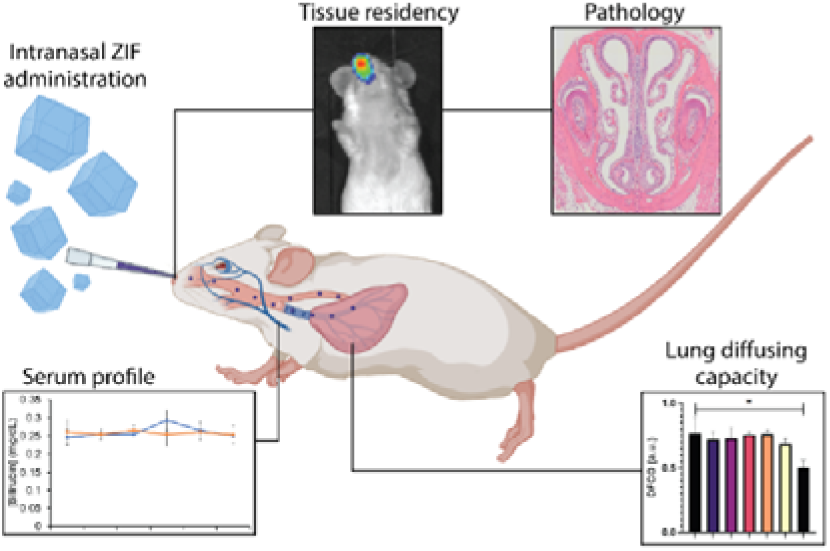

## Introduction

Vaccines are a two-hundred-year-old technology that, despite new preparations and novel formulations, still require stable low-temperature storage conditions and delivery via needle and syringe.[1, 2] Injections are one of the most frequently cited issues for poor compliance with vaccines; yet, developing alternatives is complicated because vaccines are made mainly of delicate biomolecules or lipid nanoparticles. Further, recent work has found that alternative delivery approaches can promote more robust protective immune responses, particularly in diseases that affect mucosal tissues. For instance, Forsyth *et al*. compared different UTI vaccine candidates through several delivery methods and found intranasally vaccinated groups to have the lowest bacterial load post-challenge.[3] The potential for exploration in this field is immense; not only can varying the delivery route help exploit mucosal immunity for various diseases, but it can also be pertinent, especially in vaccine development targeting respiratory infections like tuberculosis and respiratory syncytial virus. Xing *et al*. also demonstrate a comparative study between intranasal and intramuscular administration of a tuberculosis vaccine, and the intranasally vaccinated group had better survival rates post-challenge.[4] Another significant advantage of exploring intranasal delivery of vaccines is that it is easier to administer for non-medical personnel by being needle-free. This will help in vaccination drives where doctors and nurses are scarce, provide a pain-free experience, reduce biohazardous waste, and prevent cross-contamination through needles.

Targeting the mucosa can boost the efficacy of certain vaccines and combining these effects with a slow-release system can enhance the formulation’s overall performance. Presently, however, intranasal delivery systems that prolong antigen residence are few and far between and almost non-existent for lipid-based systems.[5, 6] Of all the methods being explored to build slow-release systems, biomimetic mineralization—encapsulation of natively folded biomacromolecules—within Metal-Organic Frameworks (MOFs) shows great promise as they protect antigen integrity along with a slow-release platform. MOFs are self-assembling, low-density coordination polymers and a variety of them have been explored for biomaterial protection in the recent past.[7-10] In particular, zeolitic imidazolate frameworks (ZIFs)—a class of MOF made from interconnected ligands bridged by zinc ions—have emerged as the frontrunner candidate amongst all MOFs owing to their ease of synthesis in biofriendly conditions, ability to grow on a wide variety of biomaterial surfaces while protecting their surface motifs, and improved thermal stability.[11, 12] ZIFs are unique as polymeric materials due to their kinetic lability; phosphate and carbonate salts, or albumin-rich environments readily pull the zinc from the framework causing it to dissolve completely, returning the original antigen to its pristine state.[13, 14] ZIF’s versatility in capturing, preserving, and releasing antigens is exceptional and has been used to encapsulate whole viruses,[15, 16] bacteria,[17] proteins,[18, 19] DNA,[20, 21] RNA,[22] yeast,[23] liposomes,[24, 25] enzymes,[26, 27] glycans,[28] and carbohydrates.[29] We have recently shown that ZIF-encapsulated formulations exhibit a depot effect, which significantly enhances the efficacy of vaccine models.[1, 30, 31] Additionally, MOF encapsulation of fragile biomaterials stabilizes them thermally, reducing the cost of vaccine transportation, which would otherwise need to be shipped and stored in -20°C (sometimes even -80°C) freezers.

While considerable progress has been made in studying the applicability of ZIF-8 for antigen protection and slow release, ZIF-8 is a relatively new platform for antigen delivery, and questions about its compatibility with mucosal surfaces still need to be answered. While some papers shed light on the biosafety of other MOFs *in vivo*,[32-34] rigorous studies assessing the biocompatibility of ZIF-8 in mammals, particularly when delivered intranasally, have yet to be done. In this paper, we evaluate the dose-dependent toxicity of ZIF-8 in mice delivered via intranasal administration. We look at its systemic effects and changes occurring in the respiratory system, along with other adjacent studies. Specifically, we show that ZIF-8 nanoparticles delivered intranasally have no long-term impact on the nasal mucosal surfaces or the respiratory system, and we find no evidence of dysfunction in major organs. Finally, we show that intranasal delivery of ZIF-coated liposomes enhances their residency in the sinuses four-fold.

## Results and Discussion

We synthesized nano-sized ZIF-8 and characterized it using scanning electron microscopy (SEM) (**Fig 1A**) and powder X-ray diffraction (PXRD) (**Fig 1C**) for dose-dependent studies of the MOF itself. Since ZIF-8 is intended to be used as a protective coating for biomaterials, we also prepared liposomes encapsulated in ZIF-8 (Lip@Z) with an encapsulation efficiency of 99% as a model. For imaging-based studies, the liposomes were filled with fluorescent cyanine-7 (Cy7) dye as cargo and were subsequently encapsulated in ZIF-8 (Lip(Cy7)@Z). The liposomes were characterized using dynamic light scattering (**Fig S1**) and transmission electron microscopy (**Fig S2**). Lip(Cy7)@Z was also characterized by SEM (**Fig 1B**), PXRD (**Fig 1C**), and epifluorescence microscopy (**Fig 1D**).

**Figure 1:**
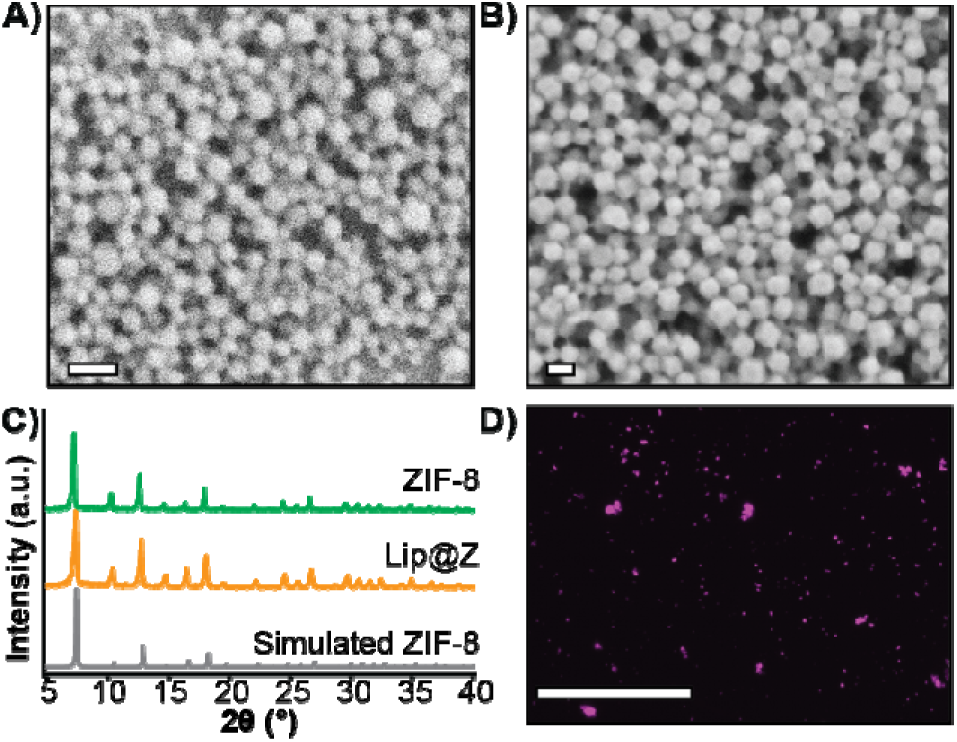
(A) SEM of ZIF-8 (scale = 200 nm), (B) SEM of Lip(Cy7)@Z (scale = 200 nm), (C) PXRD of ZIF-8 and Lip(Cy7)@Z along with simulated ZIF-8 for reference, (D) Epifluorescence micrograph of Lip(Cy7)@Z (scale = 200 µm)

Before performing *in vivo* experiments, cytotoxicity assays for ZIF-8 and Lip@Z were carried out on the A549 human lung adenocarcinoma (**Fig 2A**) and LLC Lewis lung carcinoma (**Fig S3**) cell lines. In the LLC1 model, IC_50_ of ZIF-8 was calculated as 70.52 ± 3.46 µg/mL and Lip@Z as 69.15 ± 2.54 µg/mL. In the A549 model, doses up to 300 µg/mL were relatively non-toxic, and an IC_50_ could not be established for both ZIF-8 and Lip@Z in this cell line up to 1 mg. We next assessed how long these materials would remain within the nasal cavity following intranasal administration in BALB/c mice (n = 3). The corresponding amount of encapsulated Lip(Cy7) was used as a control group (n = 3) to study the difference the ZIF-8 coating provides to tissue residency. We observed that the material cleared out of the body in under 18 h for the Lip(Cy7)@Z group, which significantly improved over the control group that lasted under 4 h (**Fig 2C**). The fluorescent intensities were plotted, and the half-life of the Lip(Cy7)@Z group was calculated to be ∼9 h, which is over four times the half-life of free liposomes (∼2.1 h) (**Fig 2B**). To assess if the zinc was bioaccumulating in major organs following dosing, *ex vivo* analysis 18 h post administration was conducted via inductively coupled plasma mass spectrometry. No significant elevation in zinc levels was observed in the liver, spleen, kidneys, and lungs compared to the liposome-only control group, indicating the zinc was likely excreted (**Fig S4**). It is worth mentioning that zinc homeostasis is highly controlled in mammals.[35]

**Figure 2:**
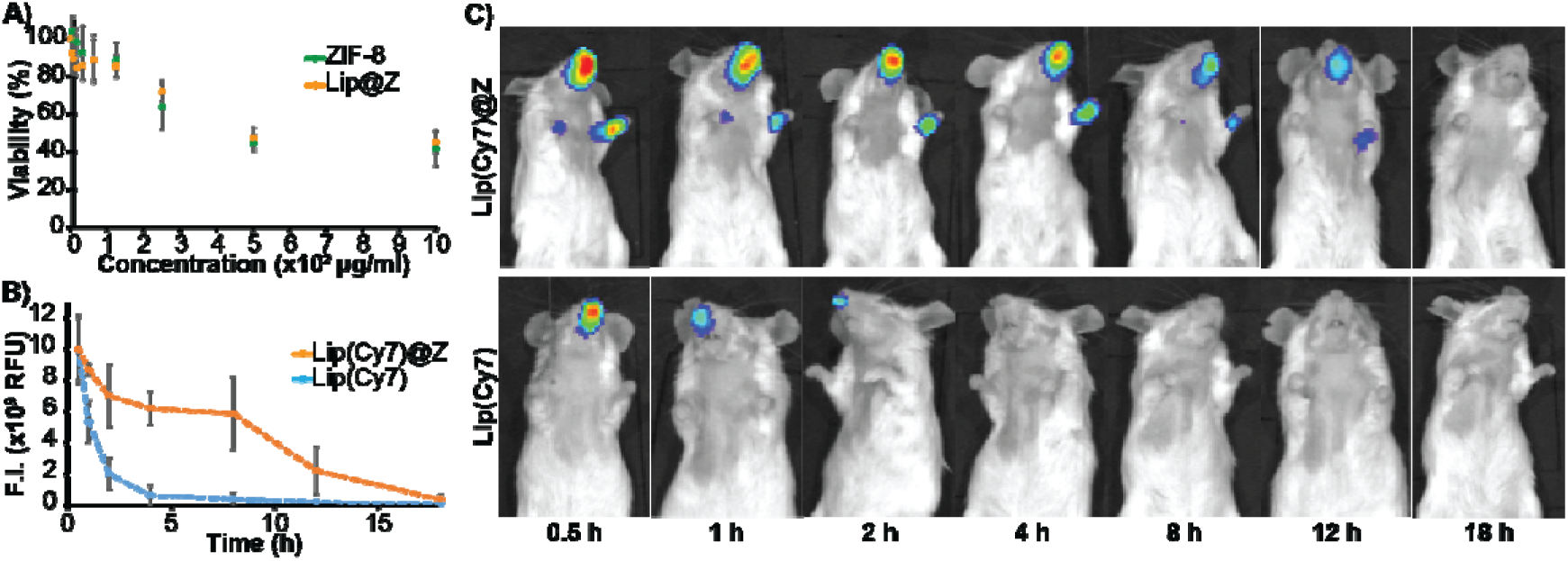
(A) Cytotoxicity of ZIF-8 and Lip@Z (4 h) in A549 cells (B) Time profile of quantified fluorescence (C) fluorescence images of mice until 18 h.

Before performing *in vivo* experiments, cytotoxicity assays for ZIF-8 and Lip@Z were carried out on the A549 adenocarcinoma (**Fig 2A**) and LLC-1 lung carcinoma cell lines (**Fig S3**). The A549 model showed a high tolerance towards the composite materials, with the IC_50_ values for ZIF-8 and Lip@Z being 433.79 ± 7.9 µg/mL and 475.72 ± 1.4 µg/mL respectively. On the more sensitive LLC-1 model, IC_50_ of ZIF-8 was calculated as 90.30 ± 6.4 µg/mL and Lip@Z as 97.39 ± 0.9 µg/mL. Next, we emulated a typical *in vivo* vaccination experiment with three doses to assess how repeated exposure to the material would affect the mucosa of the sinuses as well as liver and kidney function. ZIF-8 was administered intranasally in 4–6 week-old C57BL/6 female mice in three successive doses, each one week apart. The groups (n = 4) were divided by the amount of ZIF-8 delivered, which ranged from 50 to 1000 µg. Two more groups were added to simulate a more application-based approach. A liposome-only group (n = 4) composed of FDA-approved lipids was added as a reference nanocarrier. Lip@Z group was added as an example biocomposite material (n = 4) and administered at 1000 µg. In all cases, blood was collected submandibularly on day 1 (one day after the first dose) and day 21 (one day after the third dose) to quantify common serum proteins and enzymes to analyze whether the administered ZIF-8 has a systemic effect on their clinical chemistry. We chose to study albumin, urea, and bilirubin among the serum proteins and alkaline phosphatase (ALP), aspartate aminotransferase (AST), and γ-glutamyl transferase (GGT) as representative serum enzymes which collectively serve as biomarkers indicating kidney and liver dysfunction.[36, 37] We observed no noticeable pattern in changes in protein (**Fig 3A**) or enzyme (**Fig 3B**) levels, and any fluctuations were non-significant with respect to the saline (PBS) control group.

**Figure 3:**
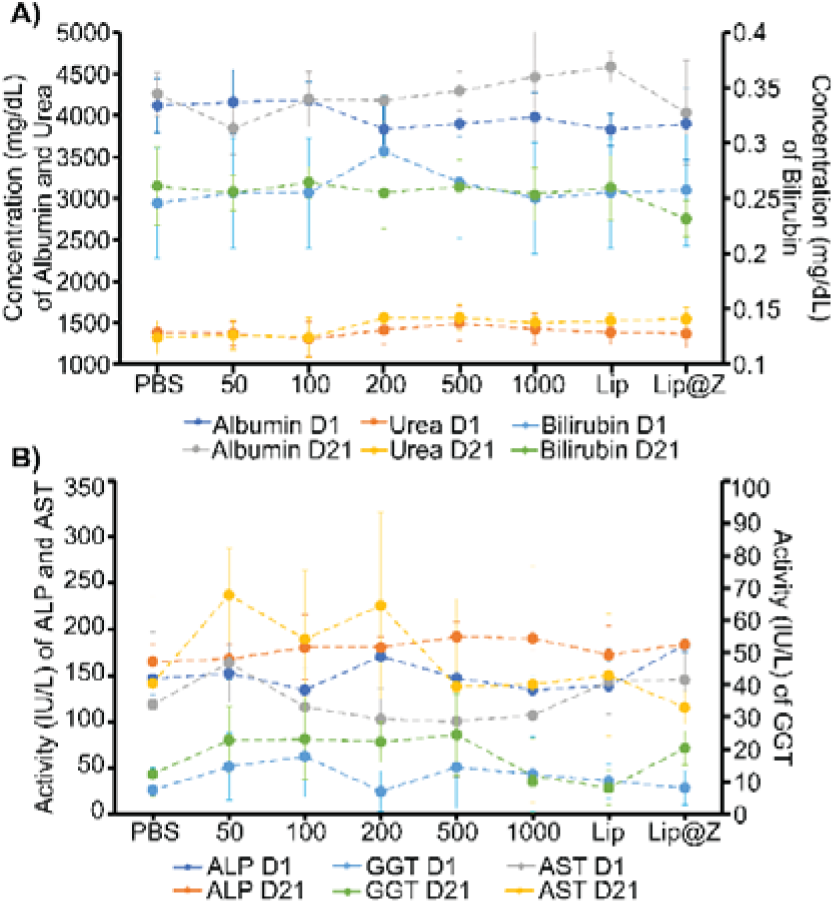
(A) Levels of albumin, urea (left axis), and bilirubin (right axis) on day 1 and day 21, (B) levels of ALP, AST (left axis), and GGT (right axis) on day 1 and day 21. On both charts, 50 to 1000 indicate the dose of ZIF-8 administered in µg.

It is important to ensure that directly administering particles into the nasal cavity does not affect the function of the respiratory system. A common test used in clinical settings is diffusing capacity of the lungs for carbon monoxide (DLCO); this test uses a gas mix containing small percentages of carbon monoxide and an inert tracer gas to measure the diffusing capacity of the lungs by the amount of carbon monoxide absorbed. This test is challenging to perform in small animal models for practical considerations. Limjunyawong *et al*. developed a close analog for this test, which measures the diffusion factor for carbon monoxide (DFCO) in mice (**Fig 4A**). [38] We used this methodology—and modified this test to be less invasive—to determine how three intranasally delivered ZIF-8 doses affect lung function. The mice from the previous experiment were used to record DFCO measurements (n = 4).

**Figure 4:**
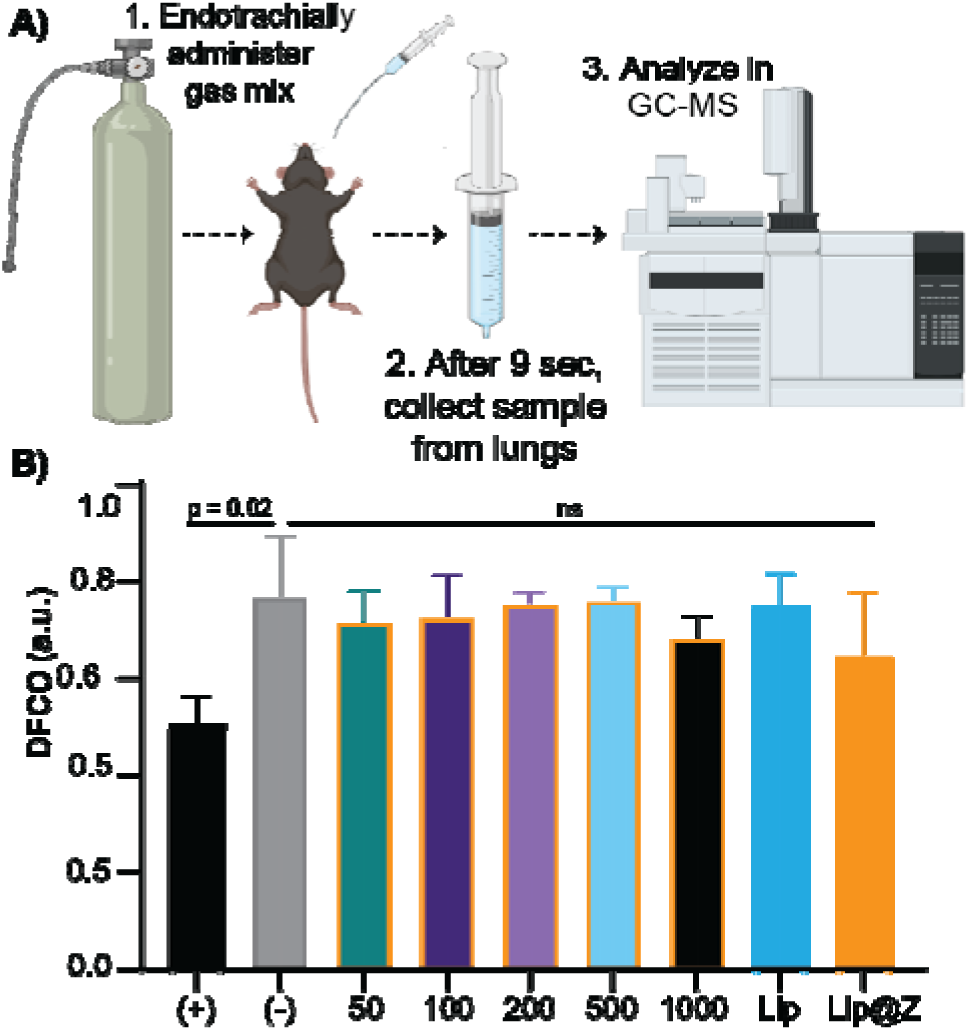
(A) Experimental scheme for DFCO test (B) DFCO measurements for all groups. (B) DFCO measurements for each group. (+) stands for positive control, mouse euthanized and used immediately after cervical dislocation, and (-) stands for the saline group. 50 to 1000 indicate the dose of ZIF-8 administered in µg. (+) and (-) are significantly different, with p = 0.02. No significant difference exists between (-) and all the ZIF, liposome, and Lip@Z groups.

All experimental groups (including Lip and Lip@Z) had similar DFCO values to that of the negative control (PBS) group, as no significance was obtained between any groups (**Fig 4B**). For positive control of damage in the respiratory system, we euthanized mice (n = 3) through isoflurane overdose followed by cervical dislocation and carried out their DFCO measurement immediately after they were euthanized. The measurement of the euthanized positive control group is significantly different compared to the negative control group. The absolute values of negative and positive control mice also correlate with the literature values.[39]

Once the mice were sacrificed after blood draws and DFCO studies, we extracted their nasal turbinates, trachea, and lungs to assess tissue damage using hematoxylin and eosin (H&E) staining. For the nasal turbinates (**Fig 5A**), we did not observe any distinct thinning of the olfactory epithelium, presence of Splendor-Hoeppli bodies, or atrophy in turbinate structure across any groups, including the 1000 µg dose. One marked difference in the groups with higher doses of ZIF-8 and the 1000 µg Lip@Z group was an increase in goblet cells, often observed after short-term exposure to a mild irritant (**Fig 5B**). However, a similar density of goblet cells was also observed in the liposome group, indicating that exposure to a variety of nanocarriers triggers mild irritation, and the issue is not specific to ZIF-8. In the lungs (**Fig 5C**), we observed little to no detachment of viable cells from the epithelial surface, no observable neutrophil infiltration in alveoli or enlarged, foamy macrophages, or any significant amount of fibrosis. There were no major pathological changes concerning the thickness of the epithelial lining or density of nuclei in the trachea across any groups (**Fig 5D**).

**Figure 5:**
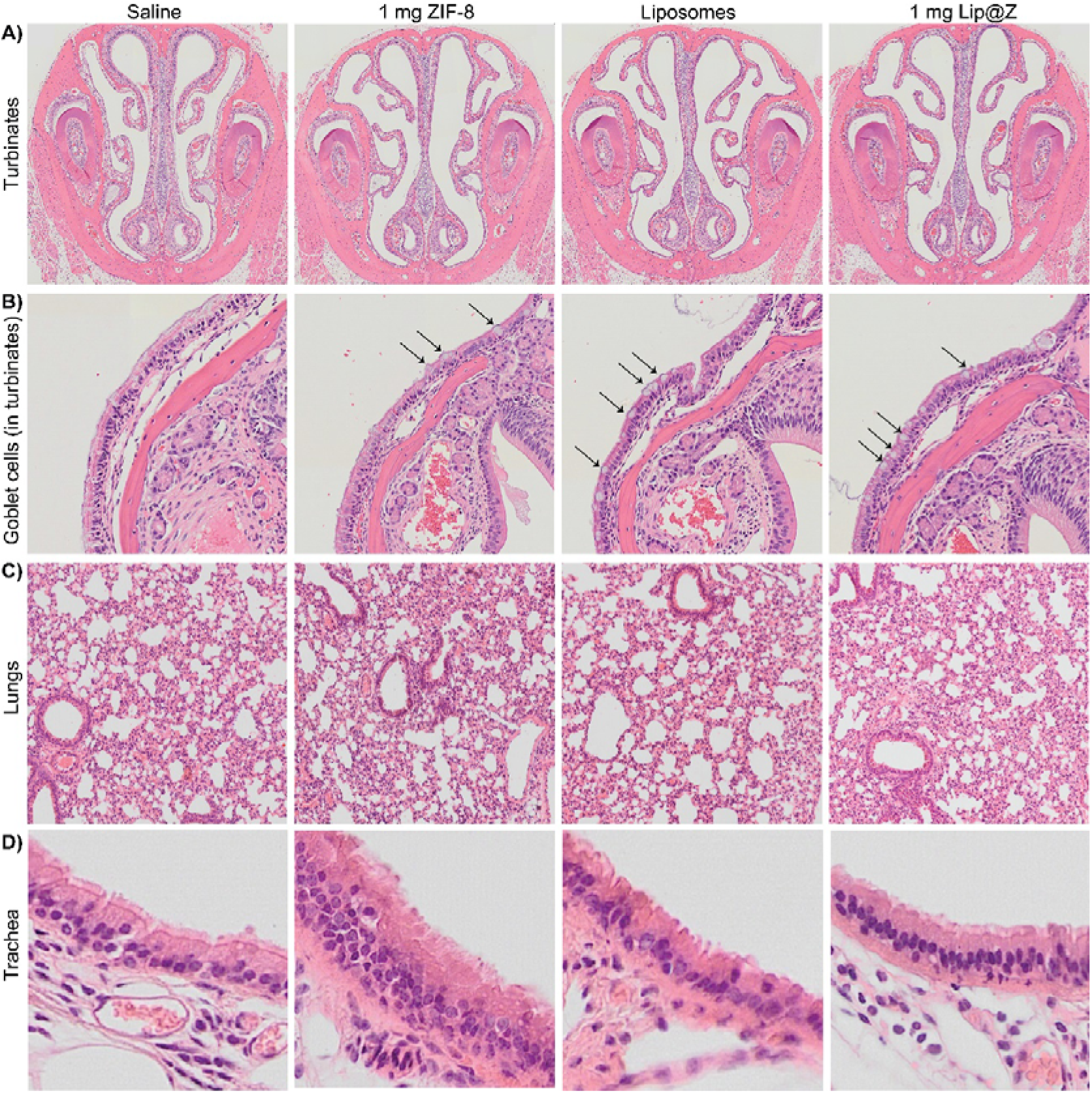
H&E for (A) turbinates, (B) arrows indicating goblet cells inside the above turbinates, (C) lungs, (D) trachea

## Conclusion

In this work, we explored the *in vivo* biosafety of ZIF-8 for application in intranasal vaccine delivery. There are various reasons why we want to encapsulate delicate biomaterials in ZIF-8. Still, if the MOF causes issues in the body, it cannot be utilized for said benefits. After investigating the systemic effects, respiratory function, and tissue pathology, we conclude that the material is safe for intranasal delivery in animal models. Additionally, we have shown that we can slow the release of materials in the nasal cavity, which can be exploited for immunological benefits when developing vaccine formulations. Having robust biosafety studies in mammals allows the scientific community to target future research to build upon this knowledge and aid in developing MOF-based intranasal vaccines to tap into mucosal immunity.

## Materials and Methods

### Materials

18:0 PC (DSPC) 1,2-distearoyl-sn-glycero-3-phosphocholine and 18:1 (Δ9-Cis) PE (DOPE) 1,2-dioleoyl-sn-glycero-3-phosphoethanolamine were purchased from Avanti Polar Lipids. MOPS 3-(4-morpholino) propane sulfonic acid, TCEP tris(2-carboxyethyl) phosphine, sodium chloride, chloroform, nitric acid (trace metal grade), hydrochloric acid, and glacial acetic acid were purchased from Thermo Fisher Scientific. Cyanine-7 was synthesized in-house; characterization has been included in the supplementary information (**Fig S5 to S11**). Zinc acetate dihydrate, 2-methylimidazole, ethanol (100%), histological grade xylenes, Harris hematoxylin solution (modified), Scott’s tap water substitute concentrate, eosin Y aqueous solution, L-glutamine solution, bromocresol green (BCG) albumin assay kit, alkaline phosphatase assay kit, aspartate aminotransferase (AST) activity assay, bilirubin assay kit, _γ_-glutamyltransferase (GGT) activity colorimetric assay kit, and urea assay kit were purchased from Millipore Sigma. HyClone phosphate buffered saline solution, HyClone Dulbecco’s modified eagle’s medium (DMEM), HyClone RPMI 1640 medium, and penicillin-streptomycin were purchased from Cytiva. FB Essence was purchased from Avantor. Cell culture grade dimethyl sulfoxide (DMSO) and histological grade ethanol (95%) were purchased from Bioworld. Deep blue cell viability kit was purchased from Biolegend. 0.3% carbon monoxide + 0.3% neon + 21% oxygen + balance nitrogen gas mix cylinder was purchased from SCI Analytical Laboratories. Isoflurane solution was purchased from Covetrus. Cytoseal 60 was purchased from Electron Microscopy Sciences.

### Cells and Animals

A549 and LLC-1 cells were received as gifts from Dr. Laurentiu Pop (Department of Radiation Oncology, UT Southwestern). Female C57BL/6 mice (4-6 weeks) and male BALB/c mice (10-12 weeks) were purchased from Charles River Laboratories. All in vivo experiments were carried out under protocol #19-06, approved by IACUC.

### Instruments

PXRD spectra were measured using a Rigaku SmartLab X-ray diffractometer. SEM micrographs were taken using Zeiss Supra 40. Liposomes were extruded using Avanti Mini Extruder and pelleted down using Sorvall MX-120 micro-ultracentrifuge. Fluorescence spectra were recorded using a Horiba Fluorolog fluorimeter. DLS measurements were carried out using Malvern Analytical Zetasizer Nano ZS. TEM micrographs were taken JEOL 1400 transmission electron microscope. Cell counting was carried out using Thermo Countess II. Colorimetric and fluorescence-based assays were analyzed using a Biotek Synergy H4 Hybrid microplate reader. Epifluorescence images were taken on EVOS FL digital inverted fluorescence microscope. Live animal imaging was done on IVIS Lumina III. Zinc concentration in organs was quantified using Agilent 7900 ICP-MS. Lung capacity gas samples were analyzed using Agilent GCMS. Tissue samples were washed on a solvent gradient using Leica ASP300 S. Paraffin embedding was carried out on HistoCore ARCADIA. Tissue samples were sliced using a Leica RM2235 microtome.

### Synthesis of ZIF, liposomes, and ZIF-encapsulated liposomes

ZIF was synthesized using zinc acetate and 2-methylimidazole, whose final concentrations in DI water were 20 mM and 1.28 M, respectively. The reaction was carried out for 3 h at RT and then washed twice. For washing, the product was centrifuged at 4000 ×g for 15 mins at RT, and the supernatant was exchanged with DI water. After washing, ZIF was dried under vacuum conditions overnight to obtain a crystalline solid. Liposomes were synthesized and encapsulated in ZIF using a previously published protocol (encapsulation with 20 mM zinc acetate and 640 mM 2-methylimidazole).[25]

### Cell culture, cytotoxicity measurement, and IC_50_ determination

A549 and LLC-1 cells were cultured in DMEM supplemented with FB essence, L-glutamine, and penicillin-streptomycin. All cells were grown in T75 flasks at 37ºC, 5% CO_2_ till 80% confluence. After detaching the cells with trypsin, trypsin was neutralized using DMEM. Cells were counted and seeded in a 96-well plate (25,000 cells/well). ZIF-8 and Lip@Z were added in concentrations ranging from 0 µg/ml to 1000 µg/ml for A549 and 0 µg/ml to 160 µg/ml for LLC-1, and incubated at 37ºC, 5% CO_2_ for 4 h. 30 min before 4 h time point, 10 μL of lysis buffer was added to appropriate wells as a negative control. At 4 h, 10 μL of the Deep blue cell viability kit dye was added to each well and incubated for 4 h. The plate was read for fluorescence at an excitation of 530 nm and an emission of 590 nm. % viability was normalized to media only and lysed cell controls and data is presented as average ± standard deviation (N = 4 with outlier analysis done in GraphPad Prism software with Grubbs’ method). IC_50_ was determined using linear regression function in excel.

### Synthesis of Cy7 dye

i. Synthesis of 2,4-dinitrophenyl p-toluenesulfonate: 2,4-Dinitrophenol (2.5 g, 13.58 mmol) was dissolved in dichloromethane (50 mL), p-toluenesulfonyl chloride (2.84 g, 14.93 mmol), and triethylamine (3.45 g, 33.95 mmol) were added successively, and the mixture was stirred for 16 h at room temperature. 50 mL water was added, and the mixture was extracted with dichloromethane (3×30 mL). The organic phase was washed with NaHCO_3_ (3×30 mL), brine (3×30 mL), and the organic fraction was evaporated under reduced pressure. The crude product was purified by trituration with hot methanol (30 mL) to give the pure product (white solid). Yield: 2.57 g (56%). ^1^H NMR (500 MHz, d6– DMSO): δ (ppm) 8.83 (d, J = 2.8 Hz, 1H), 8.69 – 8.47 (m, 1H), 7.91 – 7.71 (m, 2H), 7.68 – 7.35 (m, 3H), 2.46 (s, 3H).
ii. Synthesis of 1-(2,4-dinitrophenyl)-4-(methoxycarbonyl) pyridin-1-ium p-toluenesulfonate: A mixture of 2,4-dinitrophenyl p-toluenesulfonate (2.57 g, 2.57 mmol) and methyl isonicotinate (0.94 g, 6.90 mmol) were dissolved in toluene (7 mL/mmol). The reaction was refluxed for 16 h and left to cool to room temperature. The precipitate was filtered, washed with toluene (2 × 5 mL) and Et_2_O (2×5 mL), and dried. Yield: 0.78 g (24%). ^1^H NMR (500 MHz, DMSO) δ 9.59 (dd, J = 6.7, 1.6 Hz, 2H), 9.12 (t, J = 1.8 Hz, 1H), 8.98 (dd, J = 8.7, 2.5 Hz, 1H), 8.88 – 8.75 (m, 2H), 8.40 (d, J = 8.7 Hz, 1H), 7.45 (dd, J = 8.2, 1.9 Hz, 2H), 7.10 (d, J = 7.8 Hz, 2H), 4.05 (s, 3H), 2.29 (s, 3H).
iii. Synthesis of 4-carboxy -1-(2,4-dinitrophenyl)pyridin-1-ium p-toluenesulfonate: The carboxylic acid was prepared by hydrolysis of 1-(2,4-dinitrophenyl)-4-(methoxycarbonyl)pyridin-1-ium p-toluenesulfonate in aqueous HCl (6M, 50 mL) at 50°C for 36 h. The solvent was evaporated at reduced pressure, and the crude product was purified by recrystallization from methanol (50 mL) to give the pure product as a white solid. Yield: 0.68 g (91%). ^1^H NMR (500 MHz, DMSO) δ 9.59 (dd, J = 6.7, 1.6 Hz, 2H), 9.12 (t, J = 1.8 Hz, 1H), 8.98 (dd, J = 8.7, 2.5 Hz, 1H), 8.85 – 8.70 (m, 2H), 8.40 (d, J = 8.7 Hz, 1H), 7.45 (dd, J = 8.2, 1.9 Hz, 2H), 7.10 (d, J = 7.8 Hz, 2H), 2.29 (s, 3H).
iv. Synthesis of 3-(2,3,3-trimethyl-3H-indol-1-ium-1-yl)propane-1-sulfonate: 2,3,3-Trimethylindoline (3.00 g, 18.84 mmol) and 1,3-propane sulfone (2.30 g, 18.84 mmol) was stirred at 120°C for 1.5 h. Methanol was added to the reaction mixture and stirred at room temperature for 0.5 h. Ethyl acetate and cyclohexane were added. The white precipitate was filtered and washed with ethyl acetate. The solid was dried under reduced pressure to afford 3-(2,3,3-trimethyl-3H-indol-1-ium-1-yl)propane-1-sulfonate (4.25 g, 15.12 mmol, 80%) as a white solid. ^1^H NMR (500 MHz, MEOD): δ = 8.06−8.00 (m, 1H), 7.83−7.77 (m, 1H), 7.65−7.59 (m, 2H), 4.76 (t, J = 8.0 Hz, 2H), 3.03 (s, 2H), 2.62 (t, J = 6.6 Hz, 3H), 2.40−2.29 (m, 2H), 1.62 (s, 6H) ppm.
v. Synthesis of sodium 3-(2-((1E,3Z,5E)-4-carboxy-7-((E)-3,3-dimethyl-1-(3-sulfonatopropyl)indolin-2-ylidene)hepta-1,3,5-trien-1-yl)-3,3-dimethyl-3H-indol-1-ium-1-yl)propane-1-sulfonate: 4-Carboxy -1-(2,4-dinitrophenyl)pyridin-1-ium p-toluenesulfonate (250 mg, 0.54 mmol) and 4-bromoaniline (250 mg, 0.54 mmol) were dissolved in methanol (7 mL/mmol), and the mixture was stirred at room temperature for 0.5 h. Next, 3-(2,3-dimethyl-3H-indol-1-ium-1-yl)propane-1-sulfonate (457 mg, 1.63 mmol) and sodium acetate (265 mg, 3.24 mmol) were added, and the reaction mixture was stirred for additional 16 h at room temperature. Afterward, Et_2_O (21 mL/mmol) was added, and the mixture was placed in the freezer (−16 °C). The resulting precipitate was filtered and purified by column chromatography (silica gel, dichloromethane/methanol, 10:1, then switched to 5:1). Yield: 161 mg (45%). Deep green solid. ^1^H NMR (600 MHz, MeOD) δ 8.58 – 8.56 (m, 1H), 8.14 (d, J = 13.4 Hz, 2H), 7.84 – 7.83 (m, 1H), 7.44 (s, 1H), 7.38 (d, J = 4.1 Hz, 4H), 7.22 – 7.20 (m, 2H), 6.40 (d, J = 13.4 Hz, 4H), 4.29 – 4.26 (m, 4H), 2.97 (d, J = 6.9 Hz, 4H), 2.21 (d, J = 7.8 Hz, 4H), 1.67 (s, 12H). For C_34_H_38_N_2_NaO_8_S_2_^−^ [M – H^−^] 690.2, found 691.2.

### Tissue residency and biodistribution of intranasally-delivered ZIF

Male BALB/c mice were shaved using Nair around the lower half of their face and chest. Three groups (n = 3 each) were administered with 1 mg Lip(Cy7)@ZIF, and Lip(Cy7) made of 1 mg lipids and saline, respectively, via a micropipette. All volumes were kept under 30 µL. Animals were imaged before administration and after 30 mins, 1 h, 2 h, 4 h, 8 h, 12 h, and 18 h – which is when the fluorescence disappeared for groups. Raw data were processed using Living Image^®^ 4.7.3 software.

### Serum protein and enzyme quantification assays

Submandibular blood collection was carried out on Day 1 and Day 21 from mice. 2 µL of each serum sample was used to quantify albumin, bilirubin, urea, ALP, AST, and GGT using commercially available colorimetric kits following the manufacturer’s instructions.

### DFCO test for lung capacity

A balloon was filled with a gas mix (0.3% CO, 0.3% Ne, 21% O_2_, balance N_2_) and sealed with a rubber septum. The gas mix was collected in a 1 mL syringe and was endotracheally administered into anesthetized mice using a 14G catheter. After 9 seconds, the syringe plunger was pulled, and a gas sample was collected. The sample was injected into GCMS for analysis. DFCO was calculated using a published protocol.[39]

### Histological analysis of organs

Mice were sacrificed, and their lungs, trachea, and nasal turbinates were collected. Each organ was added to 30 ml of 4% paraformaldehyde in 1× PBS and left under shaking conditions for 48 hours. Organs were transferred to 70% ethanol in MilliQ in cassettes. Turbinates were sent for further processing to Histopathology Core, University of Texas, Southwestern. Lungs and trachea were embedded in paraffin and cooled over ice. Embedded organs were cut into 5 µm thick samples and collected on positively charged slides. Once dried, slides are dipped in [i] xylenes (10 min), [ii] xylenes (10 min), [iii] xylenes (10 min), [iv] 100% ethanol (5 min), [v] 100% ethanol (5 min), [vi] 95% ethanol (5 min), [vii] 95% ethanol (5 min), [viii] 70% ethanol (5 min), [ix] 70% ethanol (5 min), [x] 50% ethanol (10 mins), [xi] tap water (5 min), [xii] hematoxylin (2 min), [xiii] tap water (10 dunks), [xiv] tap water (10 dunks), [xv] tap water (10 dunks), [xvi] accumate (0.75 mL HCl + 300 mL 70% ethanol) (4 dunks), [xvii] tap water (10 dunks), [xviii] tap water (10 dunks), [xix] Scott’s tap water (1 min), [xx] 95% ethanol (5 min), [xxi] eosin acidified with 0.5% glacial acetic acid (1 min), [xxii] 95% ethanol (10 dunks), [xxiii] 100% ethanol (1 min), [xxiv] 100% ethanol (1 min), [xxv] xylenes (5 min), [xxvi] xylenes (5 min), [xxvii] xylenes (5 min) sequentially. Stained samples were sealed with a cover slip and imaged under a brightfield microscope.

## Acknowledgments

S. Kumari would like to thank Dr. Fang Bian of the Mass Spectrometry Core for her expertise and help with the GCMS, Jashkaran Gadhvi for arranging the H&E staining setup, and UT Southwestern Pathology Core for sectioning and initial assessment of turbinate specimens. J.J.G. acknowledges the National Science Foundation [Grant No. DMR-2003534] and the Welch Foundation [Grant No. AT-1989-20190330] for their support and funding.

## Author Contributions

S. Kumari prepared the materials, performed PXRD measurements, and conducted all *in vivo* experiments, serum assays, and DFCO studies. T.S.H. acquired SEM images and performed ICP-MS experiments. T.S.H. and R.N.E performed *in vitro* experiments and cytotoxicity studies. S. Koirala synthesized and characterized Cy7 dye. O.T. and I.T. aided with H&E. Y.H.W. acquired TEM images. S. Kumari and J.J.G. conceived the project and wrote the manuscript.

