## Supplementary material for "*In Vivo* Biocompatibility of ZIF-8 for Slow Release via Intranasal Administration": SI

**Title**

**Figure S1**

Intensity (a.u.)

| Size (nm) | Record 70: Ref Re-attempt 28Apr 3 |
| --- | --- |
| 0.4 | 0 |
| 0.463 | 0 |
| 0.536 | 0 |
| 0.621 | 0 |
| 0.719 | 0 |
| 0.833 | 0 |
| 0.965 | 0 |
| 1.12 | 0 |
| 1.29 | 0 |
| 1.5 | 0 |
| 1.74 | 0 |
| 2.01 | 0 |
| 2.33 | 0 |
| 2.7 | 0 |
| 3.12 | 0 |
| 3.62 | 0 |
| 4.19 | 0 |
| 4.85 | 0 |
| 5.61 | 0 |
| 6.5 | 0 |
| 7.53 | 0 |
| 8.72 | 0 |
| 10.1 | 0 |
| 11.7 | 0 |
| 13.5 | 0 |
| 15.7 | 0 |
| 18.2 | 0 |
| 21 | 0 |
| 24.4 | 0 |
| 28.2 | 0 |
| 32.7 | 0 |
| 37.8 | 0 |
| 43.8 | 0 |
| 50.7 | 0 |
| 58.8 | 0 |
| 68.1 | 0 |
| 78.8 | 0 |
| 91.3 | 0 |
| 106 | 0.486 |
| 122 | 3.45 |
| 142 | 8.17 |
| 164 | 13 |
| 190 | 16.2 |
| 220 | 17.1 |
| 255 | 15.5 |
| 295 | 12 |
| 342 | 7.73 |
| 396 | 3.79 |
| 459 | 1.1 |
| 531 | 0 |
| 615 | 0 |
| 712 | 0 |
| 825 | 0 |
| 955 | 0 |
| 1110 | 0 |
| 1280 | 0 |
| 1480 | 0 |
| 1720 | 0 |
| 1990 | 0 |
| 2300 | 0 |
| 2670 | 0 |
| 3090 | 0 |
| 3580 | 0 |
| 4150 | 0 |
| 4800 | 0 |
| 5560 | 0 |
| 6440 | 0 |
| 7460 | 0 |
| 8630 | 0 |

Fig S1: Size characterization of liposomes through dynamic light scattering (Average: 218.7 nm, PDI: 0.142)

**Figure S2**

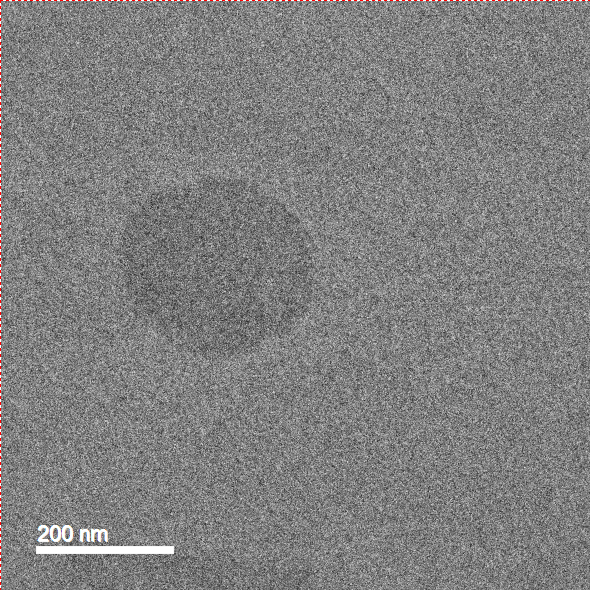

Fig S2: Transmission electron micrograph of liposome

**Figure S3**

LLC-1 Cytotoxicity

Cell Viability (%)

| Concentration (μg/ml) | 4 h ZIF-8 | 4 h Lip@Z |
| --- | --- | --- |
| 0 | 100 | 100 |
| 10 | 95.0531 | 93.9268 |
| 20 | 95.5817 | 92.6809 |
| 40 | 93.3479 | 97.1674 |
| 60 | 84.4253 | 82.5722 |
| 80 | 53.9576 | 64.2079 |
| 100 | 42.5338 | 48.1361 |
| 120 | 40.5559 | 43.3456 |
| 140 | 35.9751 | 40.5958 |
| 160 | 29.0682 | 31.8305 |

Fig S3: Cytotoxicity studies of ZIF-8 and Lip@Z on LLC-1 model

**Figure S4**

| Organ | Saline group | ZIF-8 group |
| --- | --- | --- |
| Liver | 429.6902 ± 68.41106 | 406.3947 ± 197.4786 |
| Spleen | 27.22669 ± 3.758105 | 17.48564 ± 3.534271 |
| Kidneys | 95.28645 ± 3.433586 | 73.00676 ± 20.24546 |
| Lungs | 33.89019 ± 1.768844 | 34.98579 ± 7.776183 |

Fig S4: Concentration of zinc in the liver, spleen, kidneys, and lungs in parts per billion for both the saline and ZIF-8 groups. N = 3 for each group.

**Figure S5**

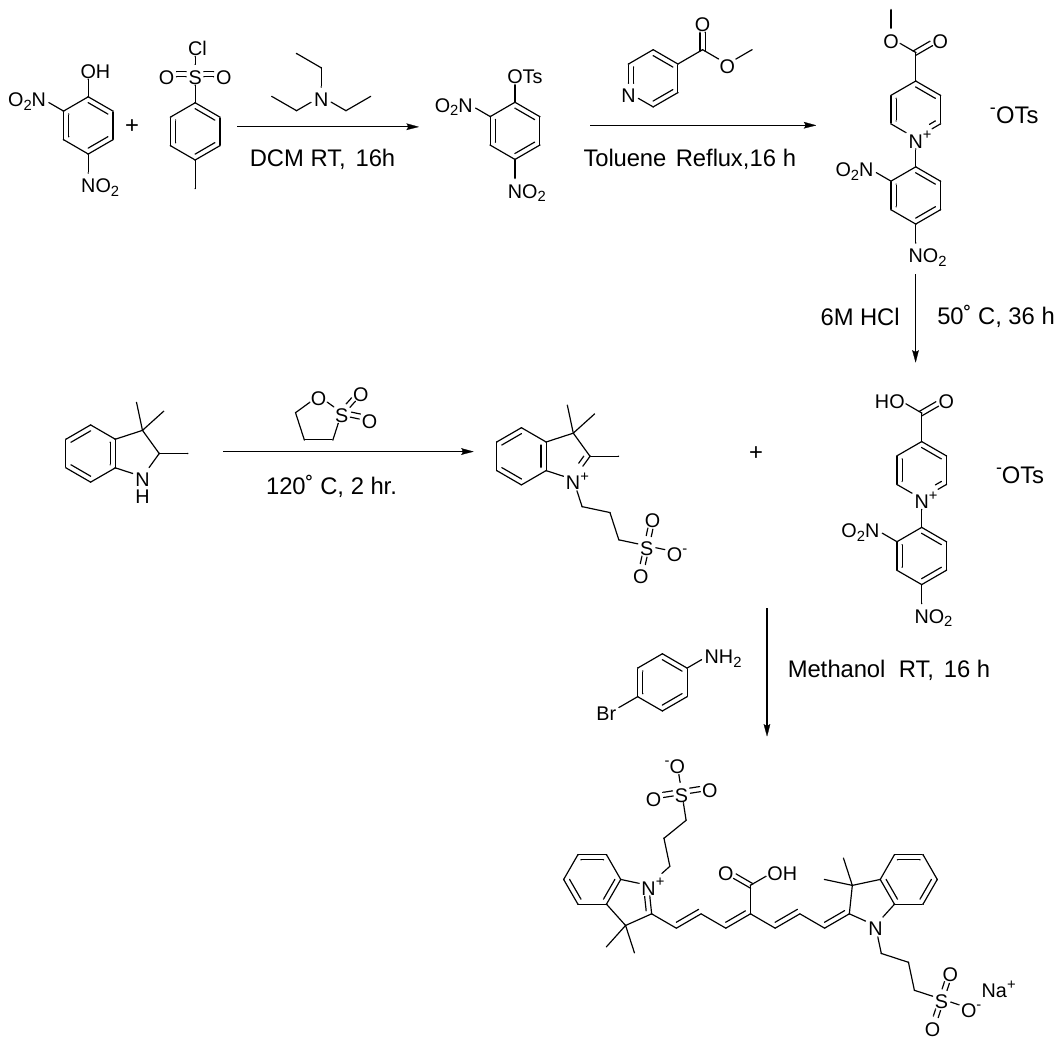

Fig S5: Scheme for synthesis of Cy7 synthesis

**Figure S6**

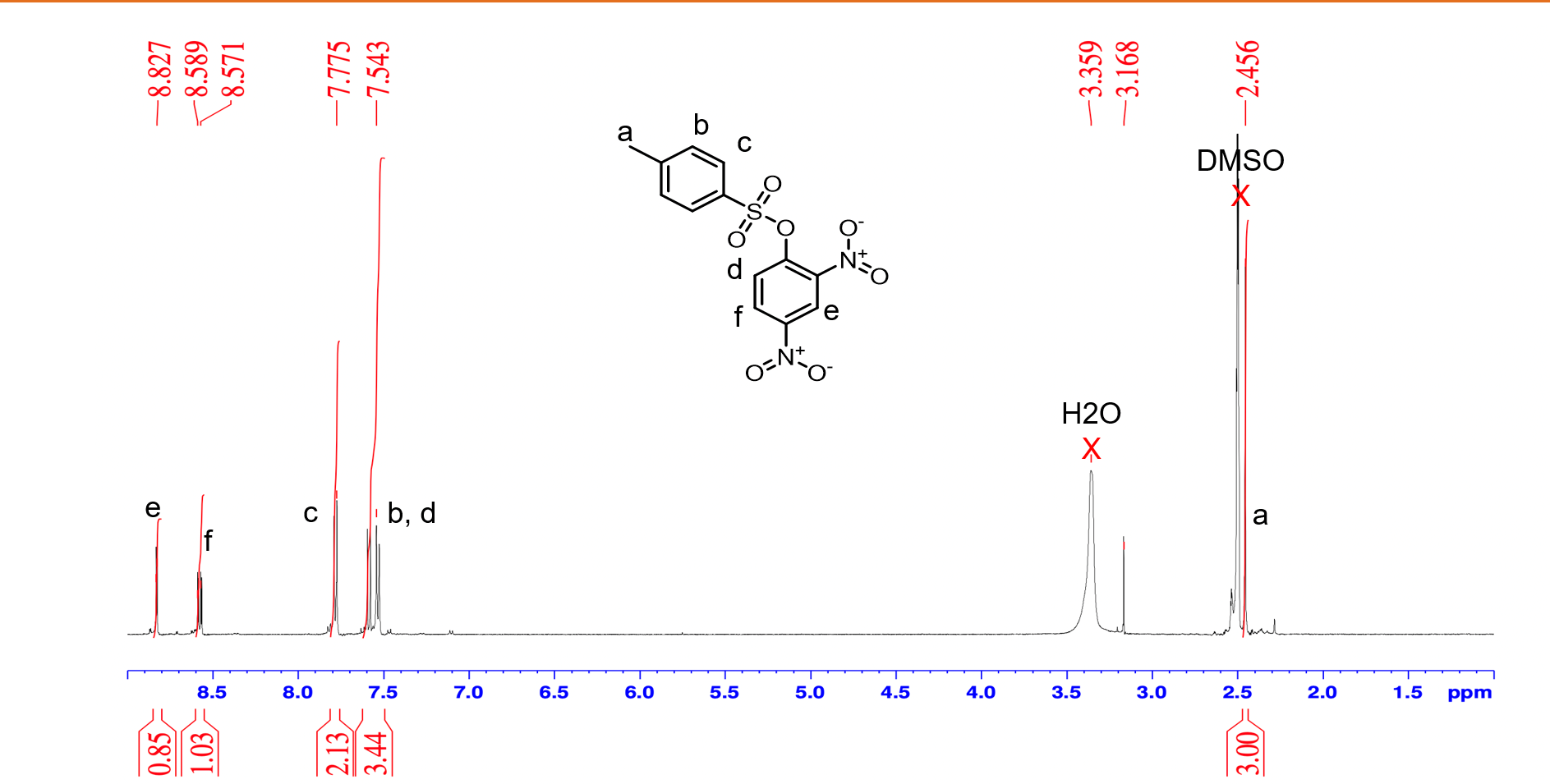

Fig S6: ^1^H NMR of 2,4-Dinitrophenyl p-toluenesulfonate

**Figure S7**

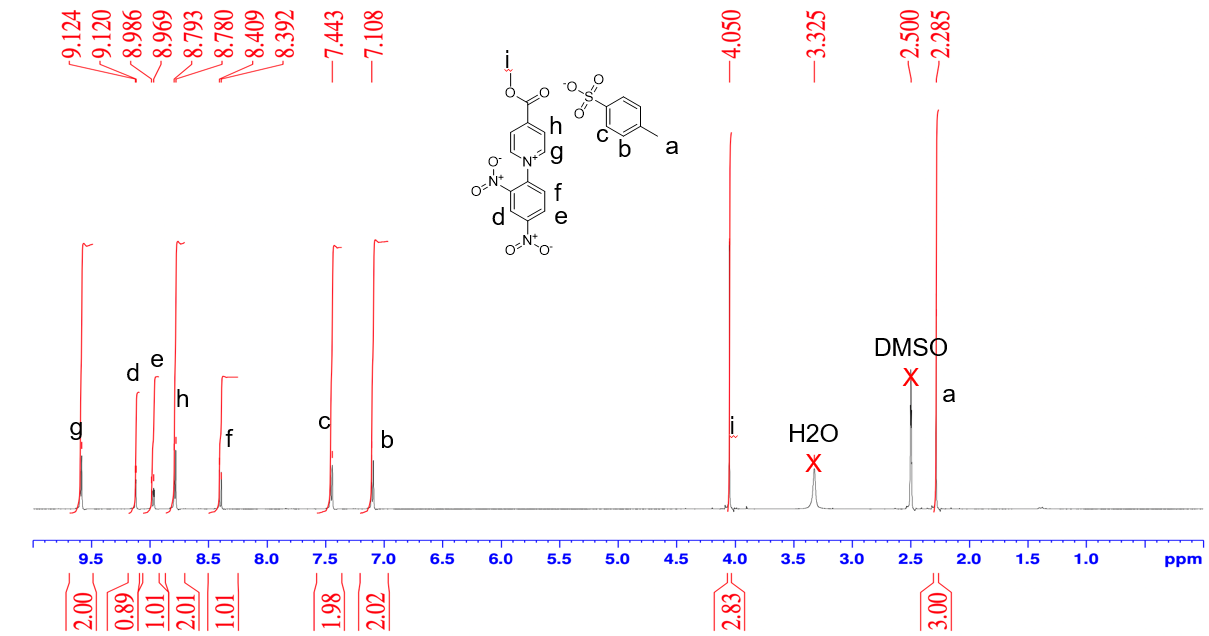

Fig S7: ^1^H NMR of 1-(2,4-dinitrophenyl)-4-(methoxycarbonyl)pyridin-1-ium p-toluenesulfonate

**Figure S8**

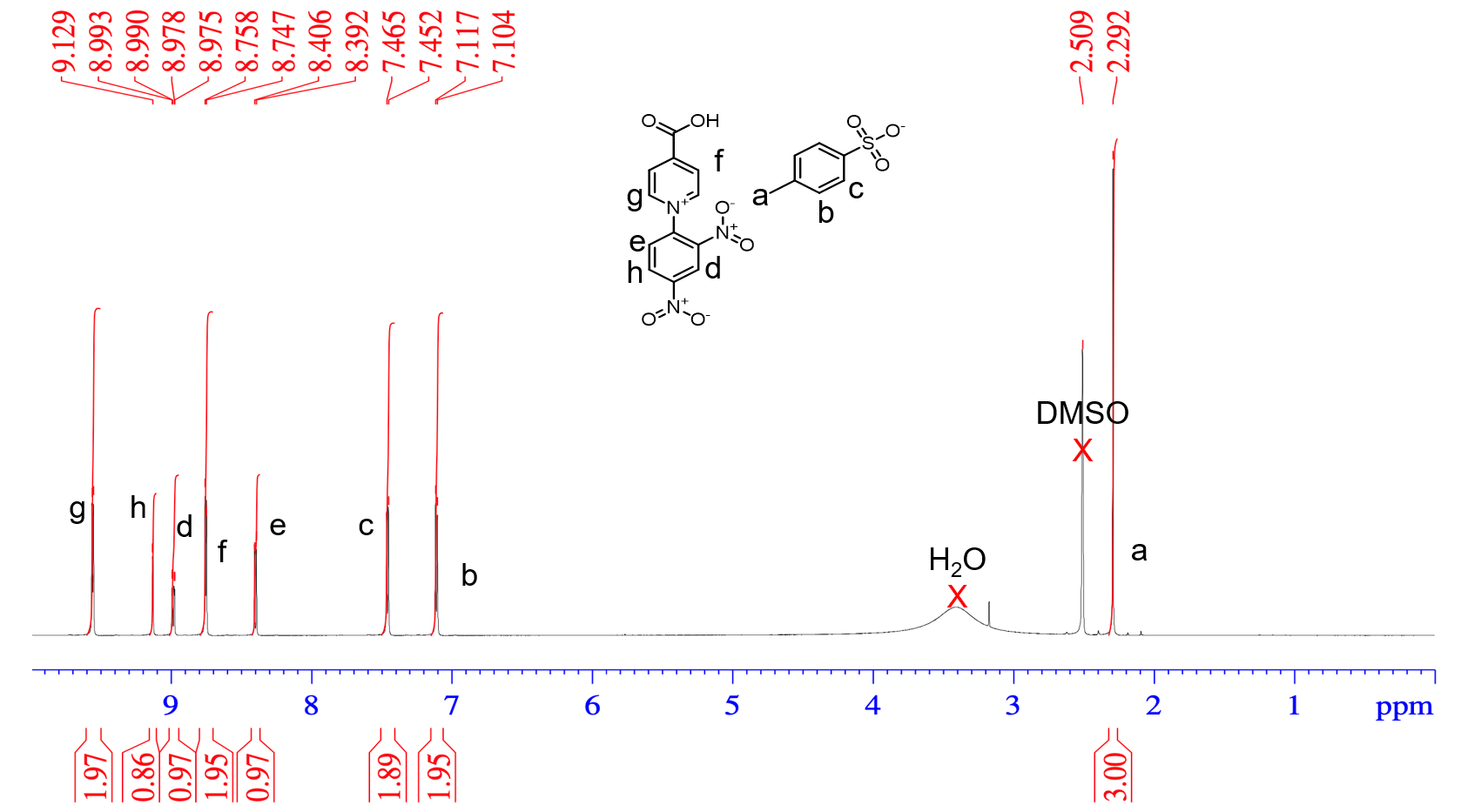

Fig S8: ^1^H NMR of 4-carboxy -1-(2,4-dinitrophenyl)pyridin-1-ium p-toluenesulfonate

**Figure S9**

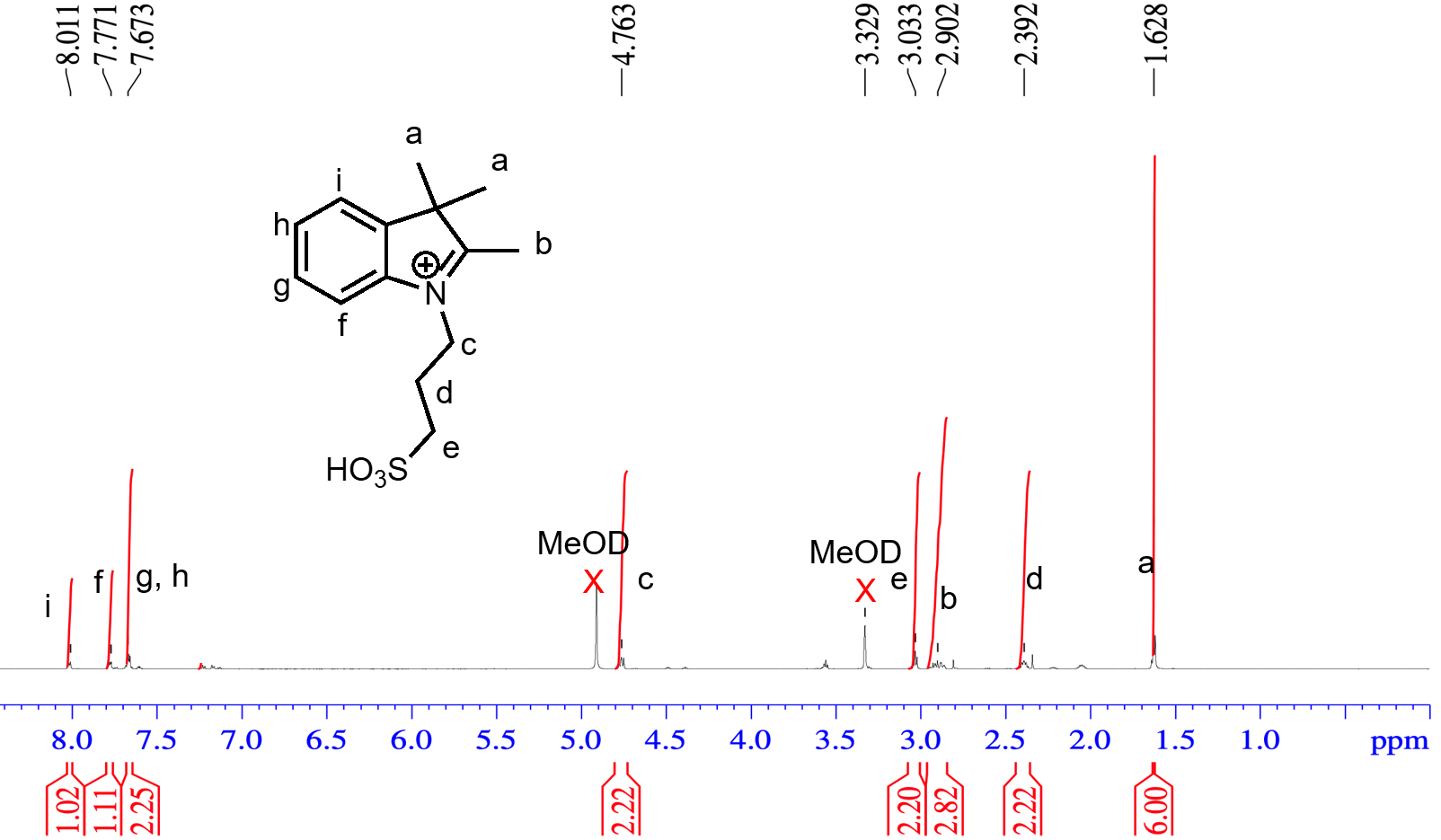

Fig S9: ^1^H NMR of 3-(2,3,3- trimethyl-3H-indol-1-ium-1-yl)propane-1-sulfonate

**Figure S10**

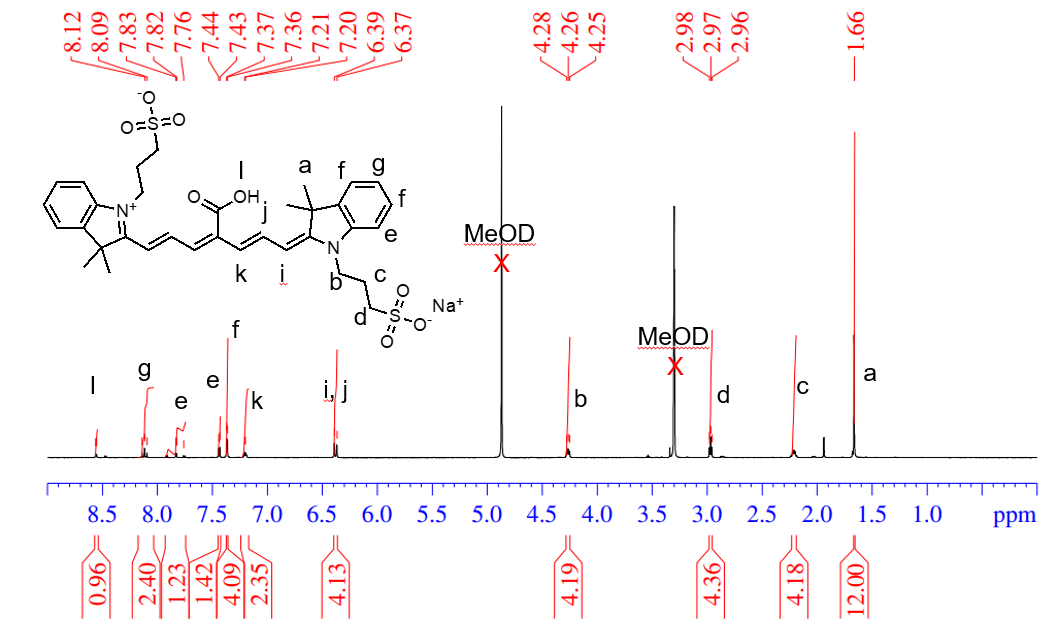

Fig S10: ^1^H NMR of sodium 3-(2-((1E,3Z,5E)-4-carboxy-7-((E)-3,3-dimethyl-1-(3-sulfonatopropyl)indolin-2- ylidene)hepta-1,3,5-trien-1-yl)-3,3-dimethyl-3H-indol-1-ium-1-yl)propane-1-sulfonate

**Figure S11**

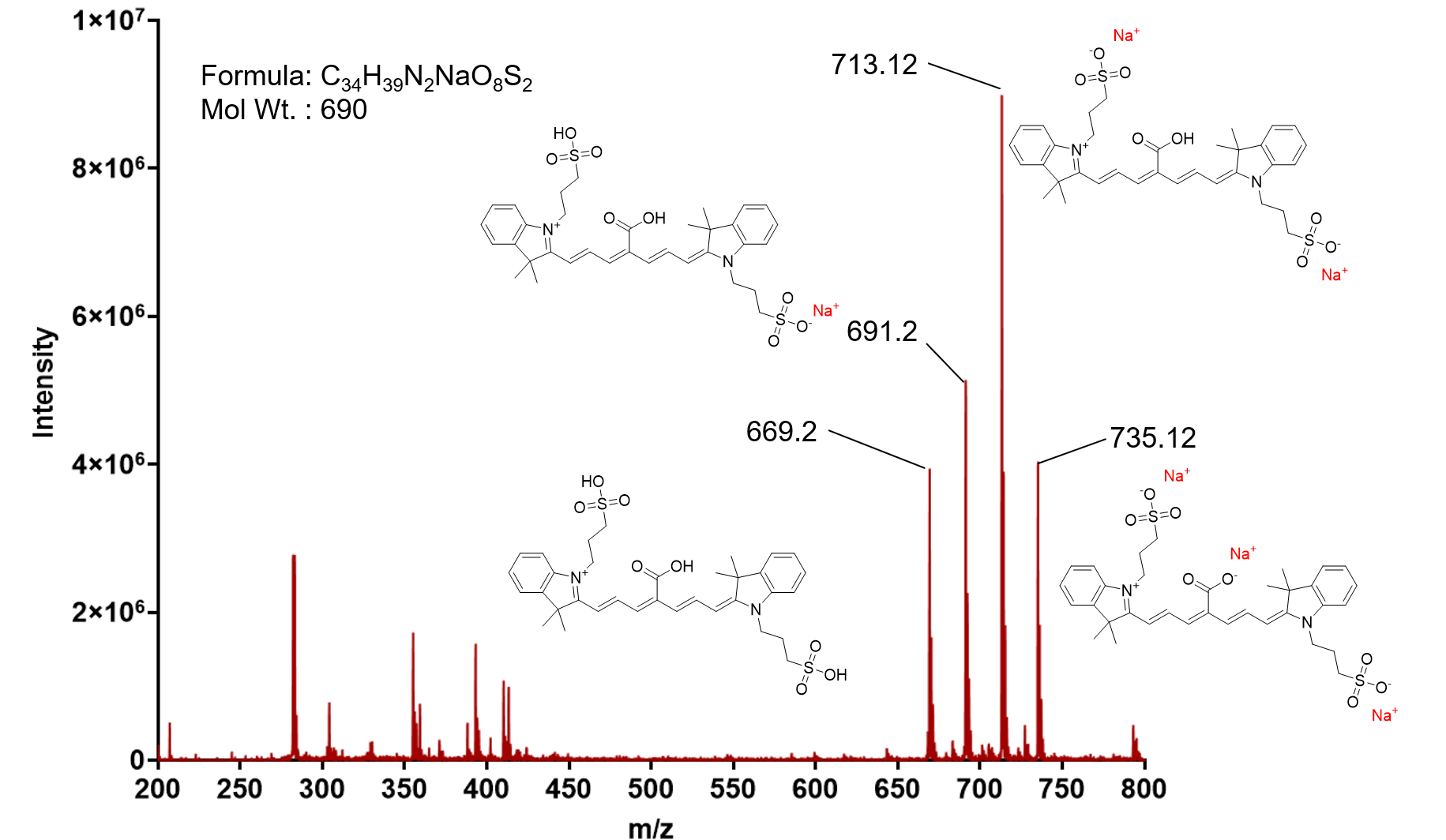

Fig S11: ESI-MS of sodium 3-(2-((1E,3Z,5E)-4-carboxy-7-((E)-3,3-dimethyl-1-(3-sulfonatopropyl)indolin-2- ylidene)hepta-1,3,5-trien-1-yl)-3,3-dimethyl-3H-indol-1-ium-1-yl)propane-1-sulfonate
